# Normative growth modeling of brain morphology reveals neuroanatomical heterogeneity and biological subtypes in children with ADHD

**DOI:** 10.1101/2024.03.16.582202

**Authors:** Xuan Bu, Yilu Zhao, Xiangyu Zheng, Zhao Fu, Kangfuxi Zhang, Xiaoyi Sun, Zaixu Cui, Mingrui Xia, Leilei Ma, Ningyu Liu, Jing Lu, Gai Zhao, Yuyin Ding, Yao Deng, Jiali Wang, Rui Chen, Haibo Zhang, Weiwei Men, Yanpei Wang, Jiahong Gao, Shuping Tan, Li Sun, Shaozheng Qin, Sha Tao, Yufeng Wang, Qi Dong, Qingjiu Cao, Li Yang, Yong He

**Affiliations:** State Key Laboratory of Cognitive Neuroscience and Learning, Beijing Normal University, Beijing, China; Peking University Sixth Hospital, Peking University Institute of Mental Health, National Clinical Research Center for Mental Disorders (Peking University Sixth Hospital), NHC Key Laboratory of Mental Health (Peking University), Beijing, China; Chinese Institute for Brain Research, Beijing, China; Center for MRI Research, Academy for Advanced Interdisciplinary Studies, Peking University, Beijing, China; Beijing City Key Laboratory for Medical Physics and Engineering, Institute of Heavy Ion Physics, School of Physics, Peking University, Beijing, China; IDG/McGovern Institute for Brain Research, Peking University, Beijing, China; Beijing Huilongguan Hospital, Peking University Huilongguan Clinical Medical School, Beijing, China; Beijing Key Laboratory of Brain Imaging and Connectomics, Beijing Normal University, Beijing, China; IDG/McGovern Institute for Brain Research, Beijing Normal University, Beijing, China

## Abstract

**Background:** Neuroimaging studies suggest substantial individual heterogeneity in brain phenotypes in attention-deficit/hyperactivity disorder (ADHD). However, how these individual-level brain phenotypes contribute to the identification of ADHD biotypes and whether these biotypes have different treatment outcomes and neurobiological underpinnings remain largely unknown.

**Methods:** We collected multisite, high-quality structural magnetic resonance imaging data from 1,006 children aged 6-14 years, including 351 children with ADHD and 655 typically developing children. Normative growth models of cortical thickness were established for 219 regions in the typically developing children. Individual-level deviations from these normal references were quantified and clustered to identify ADHD biotypes. We validated the replicability and generalizability of the ADHD biotypes using two independent datasets and evaluated the associations of the biotypes with symptomatic, cognitive, and gene expression profiles, as well as follow-up treatment outcomes.

**Findings:** No more than 10% of children with ADHD had extreme deviations in cortical thickness in a single region, suggesting high heterogeneity among individuals with ADHD. On the basis of the brain deviation maps, we discovered two robust ADHD biotypes, an infra-normal subtype with cortical thinning associated with ADHD symptoms and a supranormal subtype with cortical thickening associated with cognition. Patients with the infra-normal subtype responded better to methylphenidate than to atomoxetine, although both subtypes showed treatment efficacy. Brain deviations in the infra-normal subtype were explained by the expression levels of genes enriched in presynaptic and axonal development and polygenic risk of ADHD.

**Interpretation:** We identified anatomically distinct, clinically valuable, and biologically informed ADHD subtypes, providing insight into the neurobiological basis of clinical heterogeneity and facilitating a personalized medication strategy for ADHD patients.

**Panel: Research in context:** *Evidence before this study:* Substantial individual heterogeneity in brain phenotypes in attention-deficit/hyperactivity disorder (ADHD) motivates the need to discover homogeneous biotypes. We searched PubMed for research articles on ADHD biotypes using brain MRI published before December 1, 2023, using the search terms ((attention deficit hyperactivity disorder [Title/Abstract]) OR (ADHD [Title/Abstract])) AND ((subtypes [Title/Abstract]) OR (subgroups [Title/Abstract]) OR (subtyping [Title/Abstract])) AND ((MRI [Title/Abstract]) OR (neuroimaging [Title/Abstract]) OR (brain [Title/Abstract])) without language restrictions. Of the eight included studies, two identified ADHD biotypes using structural morphology, four used functional activity, and two used multimodal features. However, none of these studies considered the developmental effect of the brain phenotypes, examined treatment response, or investigated the genetic correlates of the biotypes.

*Added value of this study:* This study is the first to use individualized brain measures extracted from normative models to investigate ADHD biotypes in a large sample of more than 1,000 children. We identified two reproducible ADHD biotypes, characterized by distinct symptomatic, cognitive, and gene expression profiles, as well as differential treatment responses. This study advances our understanding of the neurobiological basis underlying the clinical heterogeneity of ADHD and highlights the critical need to discover ADHD biotypes using an unbiased and individualized approach.

*Implications of all the available evidence:* This study revealed remarkable neuroanatomical heterogeneity in ADHD patients and identified anatomically distinct, clinically valuable, and biologically informed ADHD biotypes. Our findings have potential value for the investigation of data-driven biotypes to evaluate treatment efficacy and facilitate personalized treatment. We also highlight the need for future studies to move beyond the understanding of ADHD solely based on the “average patient” perspective.

## Introduction

Attention-deficit/hyperactivity disorder (ADHD) is a highly prevalent and persistent neurodevelopmental disorder characterized by age-inappropriate levels of inattention, hyperactivity, and impulsivity (1). Numerous studies using clinical and psychological data have documented remarkable heterogeneity among individuals with ADHD (1, 2). Several pioneering studies have identified ADHD subtypes by dividing patients into distinct groups according to their cooccurring symptomatic, cognitive, and temperamental characteristics (3-6). However, the neurobiological substrate of this clinical heterogeneity remains elusive. The identification of ADHD biotypes could reveal their unique neurobiological mechanisms and has great potential to accelerate personalized diagnosis, prognosis, and therapeutics (7).

Neuroimaging studies have reported case-control differences in structure (8-11) and function (12, 13) between children with and without ADHD. However, most ADHD neuroimaging studies have small effect sizes and limited reproducibility. For example, two large-sample, multisite structural MRI studies reported significant (10) and nonsignificant (11) differences in cortical thickness between children with and without ADHD. These inconsistent findings reflect the heterogeneous nature of ADHD. Several recent studies have provided direct evidence for substantial heterogeneity in brain phenotypes, including cortical morphology (14) and functional connectivity (15), among ADHD individuals. These studies motivate the need to discover ADHD biotypes with homogeneous structural or functional signatures. To date, eight studies have identified ADHD biotypes using different neuroimaging measures, such as structural morphology (16, 17), functional connectivity (18-21), and combinations of both features (22, 23). These results have already begun to provide insights into the neurophysiological heterogeneity of ADHD.

Despite this substantial research effort, one limitation of existing ADHD biotype studies is their use of brain-derived phenotypes that ignore developmental effects in children with ADHD. Neuroimaging studies have suggested delayed maturation of cortical thickness (24) and functional connectivity (25) in the frontal, temporal and parietal regions in children with ADHD. Without considering development, these subtyping approaches may result in biased biotypes driven by age rather than brain characteristics per se. Furthermore, the replicability and generalizability of the ADHD biotypes across independent datasets have not been well established. Another limitation is that previous studies have not examined whether ADHD biotypes are associated with distinct treatment outcomes or unique molecular mechanisms, such as gene expression signatures. Establishing replicable and clinically and biologically valuable ADHD subtypes remains a challenge.

To address this challenge, we collected multisite, high-quality structural MRI data from a large sample of 1,006 children aged 6-14 years. An overview of the study workflow is shown in Figure 1. We first established normative growth models of cortical thickness in 219 brain regions to characterize the neuroanatomical heterogeneity of ADHD. The cortical thickness signature was used for normative modeling since it reflects the neural cytoarchitecture and is sensitive to cortical development (26). Normative modeling (15, 27) is a valuable statistical approach that relates demographic characteristics (e.g., age and sex) to brain phenotypes in a normal population and quantifies individual-level deviations from the normal reference. We then estimated individual deviations in cortical thickness in ADHD patients and applied spectral clustering analysis to identify homogeneous subtypes. We validated the replicability and generalizability of the ADHD biotypes using two independent datasets and evaluated the associations of the biotypes with cognitive and symptomatic measures via multivariate partial least squares (PLS) regression. Finally, we examined whether and how ADHD biotypes are related to different treatment responses and gene expression profiles.

**Figure 1.**
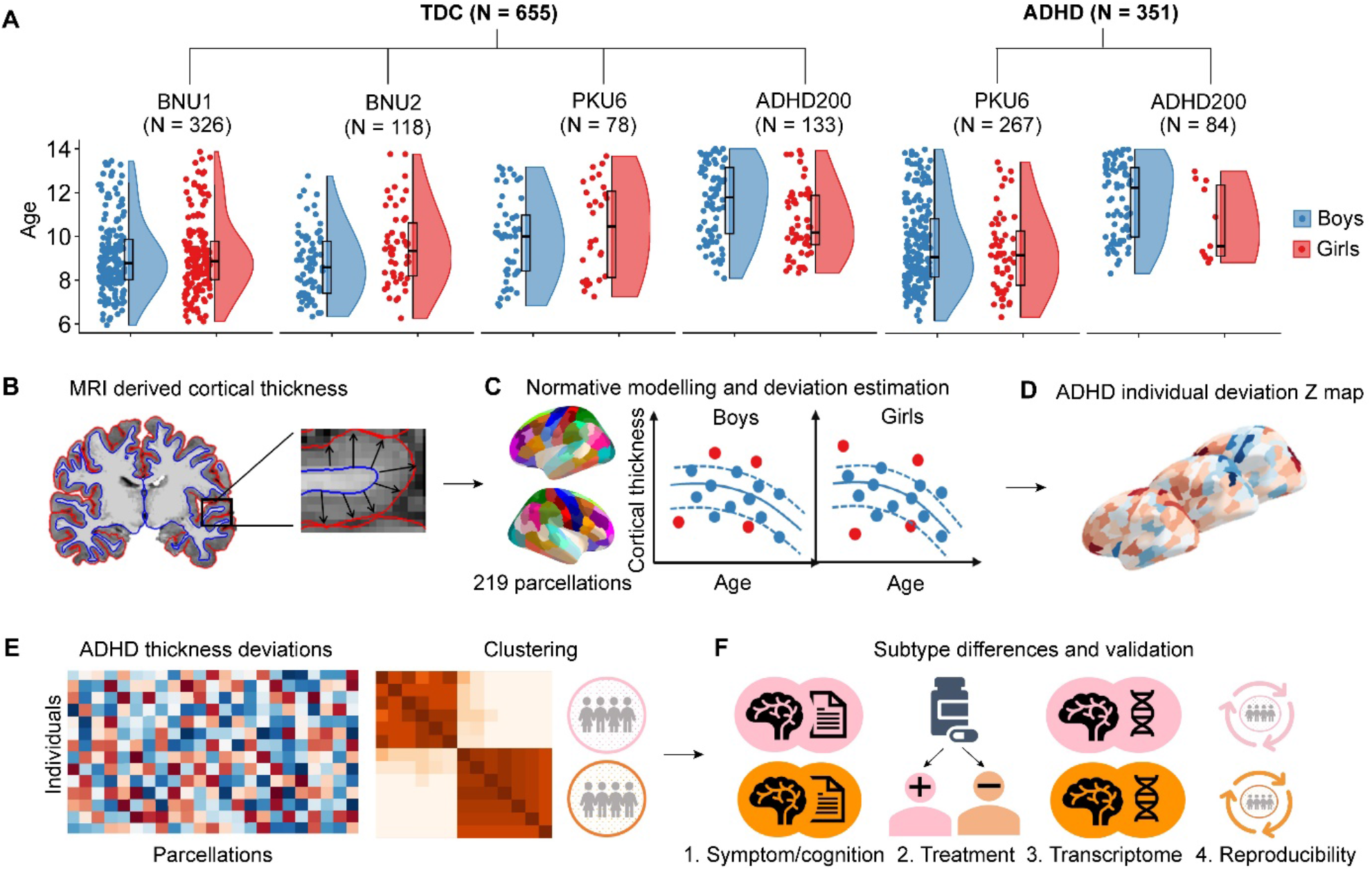
Study demographics and schematic diagram of methodology. **(A)** Box and violin plots represent the age and sex distribution of each cohort. **(B)** Regional cortical thickness was calculated for each participant. **(C)** Sex-specific normative models were estimated for each cortical parcellation. For a given individual with ADHD, the cortical thickness value of each region was compared with the corresponding normative models. **(D)** The brain maps show the cortical thickness deviation profile of each ADHD individual, representing individual differences across the whole brain. **(E)** All ADHD individuals were grouped into different brain-based subtypes using a clustering analysis based on individual cortical thickness deviations. **(F)** Subtype differences in symptom/cognition, treatment, transcriptome, and subtype reproducibility were evaluated.

## Methods

### Participants

The present study included three independent neuroimaging datasets from China: the Children Brain Development Project (BNU cohort), the Peking University Sixth Hospital (PKU6 cohort), and the publicly available ADHD200 Study (ADHD200 cohort) (28). After application of rigorous quality control (Supplementary Methods), our final sample included a total of 1,006 participants, with 655 typically developing children (TDCs) (age: 6.0-14.0 years; 369 boys; BNU, PKU6, and ADHD200) and 351 children with ADHD (age: 6.2-14.0 years; 288 boys; PKU6 and ADHD200) (Figure 1A, Table 1). This study was approved by the Ethics Committee of Beijing Normal University (BNU cohort) and Peking University Sixth Hospital (PKU6 and ADHD200 cohorts), and written informed consent was obtained from all participants and their parents.

**Table 1.** Demographics for participants.

|  | TDC |  |  |  | ADHD |  | Total |  |
| --- | --- | --- | --- | --- | --- | --- | --- | --- |
|  | BNU1 | BNU2 | PKU6 | ADHD200 | PKU6 | ADHD200 | ADHD | TDC |
| **Sample size** | 326 | 118 | 78 | 133 | 267 | 84 | 351 | 655 |
| **Demographics** |  |  |  |  |  |  |  |  |
| Age range | 6.0-13.9 | 6.0-13.9 | 6.8-13.7 | 8.1-14.0 | 6.2-14.0 | 8.3-14.0 | 6.2-14.0 | 6.0-14.0 |
| Age, mean ± SD | 9.16 (1.70) | 9.22 (1.67) | 10.06 (1.95) | 11.19 (1.70) | 9.53 (1.91) | 11.55 (1.73) | 10.02 (2.05) | 9.67 (1.90) |
| Sex, n Males | 173 (53%) | 68 (58%) | 50 (64%) | 78 (59%) | 215 (81%) | 73 (87%) | 288 (82%) | 369 (56%) |
| IQ, mean ± SD | .. | .. | 113.41 (14.66) | 118.14 (13.07) | 105.50 (16.63) | 107.07 (13.21) | 105.95 (15.72) | 116.56 (13.76) |
| **Symptoms** |  |  |  |  |  |  |  |  |
| Inattention | .. | .. | .. | .. | 26.01 (4.05) | 27.85 (4.12) | 26.48 (4.17) | .. |
| Hyperactivity-Impulsivity | .. | .. | .. | .. | 19.98 (5.84) | 22.50 (6.44) | 20.60 (6.21) | .. |
| Total | .. | .. | .. | .. | 46.00 (8.19) | 50.21 (8.00) | 47.05 (8.51) | .. |
| **Comorbidities** |  |  |  |  |  |  |  |  |
| ODD | .. | .. | .. | .. | 7 (3%) | 25 (30%) | 32 (9%) | .. |
| CD | .. | .. | .. | .. | 1 (0.4%) | 2 (2%) | 3 (0.9%) | .. |
| LD | .. | .. | .. | .. | 0 (0%) | 10 (12%) | 10 (3%) | .. |
| Tics | .. | .. | .. | .. | 3 (1%) | 7 (8%) | 10 (3%) | .. |
| Others* | .. | .. | .. | .. | 0 (0%) | 3 (4%) | 3 (0.9%) | .. |
| **Medication history** | .. | .. | .. | .. | 0 (0%) | 26 (31%) | 26 (7) | .. |

Data are mean (SD) or n (%). ADHD200 cohort only included PKU site. TDC, typically developing controls; ADHD, attention-deficit/hyperactivity disorder; ODD: oppositional defiance disorder; CD: conduct disorder; LD: learning disability. \*Others: social anxiety disorder, depression, social phobia.

### Image acquisition and calculation of cortical thickness

High-resolution T1-weighted structural MRI data were obtained for each dataset using 3.0 T MRI scanners. For each individual, the pial surface and gray matter/white matter boundary surfaces were reconstructed using FreeSurfer (v 6.0). Cortical thickness was measured as the distance between linked vertices in the pial and gray matter/white matter surfaces. Using a priori Desikan–Killiany atlas (29, 30), the mean cortical thickness was calculated for 219 regions. For details, see Supplementary Methods.

### Constructing normative growth models of cortical morphology

Using cortical thickness data from TDCs (n = 655, age range: 6.0-14.0 years), we created normative growth models of 219 cortical regions using the PCNtoolkit (31). Using hierarchical Bayesian regression (HBR), we modeled cortical thickness as a function of age, sex, and site and estimated the normative variance in thickness and the uncertainty of prediction. HBR allows for both linear and nonlinear modeling and shows superiority in addressing site effects in model estimation (32, 33). Model generalization was assessed by 10-fold cross-validation, with the model fit for each region measured using standardized mean squared error, mean standardized log-loss, and explained variance (Supplementary Methods).

### Identifying ADHD biotypes using spectral clustering

For each individual in the ADHD group (n = 351), the cortical thickness map was positioned onto the normative model to estimate individual deviations. For a given child with ADHD, *i*, the deviation score *Z* of the region, *j*, was calculated as follows:

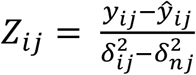

where *y*_*ij*_ is the true cortical thickness, *ŷ*_*ij*_ is the predicted cortical thickness, 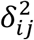 is the prediction uncertainty, and 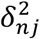 is the normative variance. Extreme positive or negative deviations were defined as |*Z*| *>* 2 (34, 35). Thus, we acquired 219 brain deviations for each child with ADHD.

To identify ADHD biotypes, we performed spectral clustering analysis on 351 ADHD deviation maps using the SNFtool (36). Briefly, the interparticipant similarity matrix (351×351) was estimated by calculating the Euclidean distance between the brain deviation profiles of every pair of participants. Spectral clustering was subsequently applied to the similarity matrix to cluster the participants into subgroups. The optimal number of clusters was determined according to eigengaps and rotation costs. Bootstrapping (1,000 iterations, 80% of 351 samples) was employed to validate the robustness of the subtyping. To evaluate the reproducibility of the ADHD biotypes, we repeated the above analyses within each site and assessed the similarity of the results between sites (Supplementary Methods).

We also compared cortical thickness differences between ADHD biotypes and TDCs using analysis of covariance (ANCOVA) with age, sex, and medication as covariates. The ComBat approach (37) was used to correct for site effects. Multiple comparisons were corrected using the false discovery rate (FDR) at *q* < 0.05.

### Relationships between cortical deviation signatures and demographic characteristics, symptoms, and cognition in ADHD biotypes

To examine whether the ADHD biotypes exhibited differences in demographic characteristics (sex and age), symptoms (inattention, hyperactivity-impulsivity, and total score), and cognitive function (delay aversion, sustained attention, response inhibition, spatial working memory, spatial planning, attention flexibility), we performed both univariate and multivariate analyses. The chi-square test was used to evaluate biotype differences in sex. General linear models were applied for assessing subtype differences in symptoms and cognitive function, with age, sex, site, and medication as covariates. We then used multivariate PLS regression for each biotype to explore the relationship between brain deviations (predictor variables) and symptoms/cognition (response variables). The statistical significance of the PLS components was evaluated using permutation tests (n = 5,000). For the significant PLS components, we calculated Pearson correlations between the brain score and symptom/cognition score, and the significance of the correlations was determined by permutation tests (n = 5,000). To assess the loading stability, we applied bootstrapping resampling (n = 5,000) to calculate the Z score. For details, see Supplementary Methods.

### ADHD biotype differences in treatment response

We further investigated whether the identified ADHD subtypes responded differently to treatment. In the PKU6 cohort, 102 children with ADHD underwent 12 weeks of medication treatment with either methylphenidate or atomoxetine, and their symptoms were assessed both before and after treatment. We examined ADHD biotype differences in treatment response using a three-way (drug type, treatment, and subtype) repeated-measures analysis of variance (ANOVA), with each symptom score as the dependent variable. If there was a three-way interaction, post hoc tests were performed within each subtype to examine the drug type by treatment interaction and the main effects of drug type and treatment.

### Transcriptomic profiling of biotype-specific changes in cortical thickness

PLS analysis was used to evaluate the associations between biotype-specific changes in cortical thickness (*T*-map) and gene expression profiles. Human brain microarray samples were sourced from the Allen Human Brain Atlas (AHBA) (n = 6 donors) (38). After preprocessing using the abagen toolbox (39), the expression profiles of 8,564 genes across 111 cortical regions were obtained. The statistical significance of the PLS components was tested using spin-based spatial permutation tests (40, 41) (n = 5,000). For significant PLS components, the Pearson correlation coefficient between the gene score and brain score was calculated, and the statistical significance was tested using spin tests (n = 5,000). We transformed the gene weight into a *Z* score by dividing by the standard error estimated from bootstrapping (n = 5,000) (42). Using univariate one-sample *Z* tests, significant genes (FDR-corrected, q < 0.05) were identified and ranked separately as positive or negative weighted genes. For details, see Supplementary Methods.

To determine the biological implications of the identified genes, we performed Gene Ontology (GO) enrichment analysis using Metascape (43) (https://metascape.org). We selected three ontology categories, namely, biological process, molecular function, and cellular component. The significantly ranked positive and negative genes were subjected to separate analyses. To alleviate redundancy in GO terms, Metascape hierarchically clustered all the significant terms into clusters of similar terms based on similarity with the kappa test score. A threshold kappa score of 0.3 was applied to split the tree into separate clusters. The most significant terms (lowest p value) within each cluster were used to represent the cluster. Finally, we investigated whether the PLS gene lists shared enrichment with risk genes for ADHD that were derived from recent GWAS studies (44-48). We performed a multigene list meta-analysis (43) between the PLS-positive (PLS+) and PLS-negative (PLS-) gene lists and the ADHD GWAS gene list, respectively, to determine whether there were shared enrichment terms or enrichment terms selectively attributed to specific gene lists. The significance threshold for enrichment was set at FDR *q* < 0.05.

## Results

### Normative growth models of cortical thickness reveal remarkable neuroanatomical heterogeneity in ADHD

We constructed normative growth models of cortical thickness for 219 brain regions (for model performance, see Figure S1). Both boys and girls showed cortical thinning with age in most regions. On the basis of these growth models, we found that no more than 10% of the children with ADHD displayed extreme deviations in a single brain region (Figure 2A), and more than 80% of the children with ADHD had at least one brain region with extreme positive or negative deviation (|*Z*| *>* 2) (Figure 2A). These results suggest substantial heterogeneity in cortical thickness signatures among children with ADHD.

**Figure 2.**
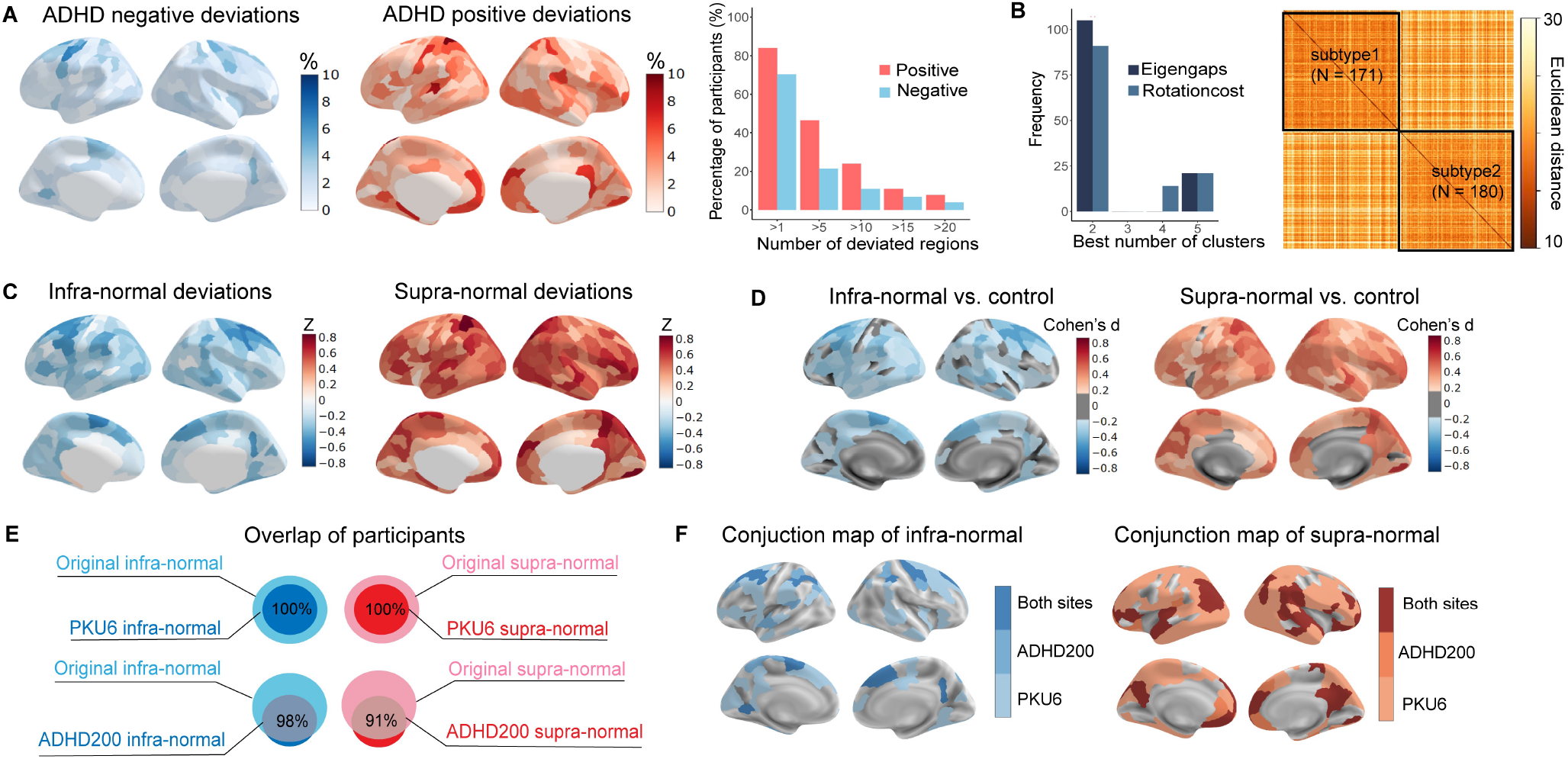
Normative models of cortical thickness revealed heterogeneity in ADHD and two ADHD subtypes were identified. **(A)** Spatial distribution map shows the percentage of ADHD children with extreme deviation (|Z| < 2) in each region. Bar plot depicts the distribution of the number of extreme positive (red) and negative (blue) deviated regions per ADHD child. **(B)** Determination of the best number of ADHD subtypes (left) and the similarity of the cortical thickness deviation patterns between participants (right). **(C)** Average cortical thickness deviation in the infra-normal subtype (left) and the supra-normal subtype (right). **(D)** Cortical thickness differences between each ADHD biotype and controls (left: infra-normal subtype vs. controls, right: supra-normal subtype vs. controls), which were identified by ANCOVA analysis. **(E)** Venn diagram depicting the degree of overlap between the subject IDs of the single-site clustering and those of the original biotypes. **(F)** Case-control differences in cortical thickness were obtained for each biotype (left: infra-normal subtype vs. controls, right: supra-normal subtype vs. controls) at each site. The conjunction maps shown here indicate a high reproducibility of results between two independent sites. Dark colors denote overlapping regions between two sites and light colors denote site-specific regions.

### Two robust ADHD biotypes characterized by distinct brain deviation profiles

Using spectral clustering analysis, we identified two ADHD biotypes with unique patterns of cortical thickness deviations (Figure 2B, Figure S2). Specifically, subtype 1, termed the infranormal subtype, exhibited widespread negative deviations, primarily in the superior and middle frontal, precentral, and superior parietal cortices (N = 171; boys = 139; Figure 2C, Figure S3A), whereas subtype 2, termed the supra-normal subtype, exhibited widespread positive deviations, primarily in the superior parietal, precuneus, medial and lateral prefrontal, and lateral temporal cortices (N = 180; boys = 149; Figure 2C, Figure S3B).

According to our case-control comparison (Figure S4), infra-normal subtype exhibited cortical thinning across widespread regions compared to TDCs (|Cohen’s d|: 0.17-0.49, p_FDR_ < 0.05; Figure 2D). Conversely, supra-normal subtype exhibited cortical thickening in extensive regions (|Cohen’s d|: 0.17-0.62, p_FDR_ < 0.05; Figure 2D). Importantly, no significant difference in cortical thickness was observed between all ADHD patients and controls.

We also identified two ADHD biotypes within each site. In the PKU6 cohort, the subject ID of each subtype had 100% overlap with that of the original subtype (Figure 2E). In the ADHD200 cohort, the subject ID of infra-normal subtype had 98% overlap with that of the original infranormal subtype, and the subject ID of supra-normal subtype had 91% overlap with that of the supra-normal subtype (Figure 2E). Subtype-specific alterations in cortical thickness also showed similar patterns between two sites (Figure 2F, Figure S5). These results suggest the high replicability and generalizability of ADHD biotypes across independent datasets.

### Subtype-related brain deviations relating to symptoms and cognition

According to univariate analysis, the two ADHD biotypes did not differ in age, sex, symptoms, or cognitive functions (Table S1). However, PLS regression revealed that the two subtypes had distinct associations between brain deviations and symptomatic and cognitive profiles.

In the infra-normal biotype, the first component of the PLS model explained 35.7% of the variance in ADHD symptoms (p_permutation_ = 0.038; Figure 3A), with a significant correlation between brain scores and symptom scores (r = 0.68, p_permutation_ < 0.001; Figure 3B). All three symptomatic variables contributed to this association (Figure 3C, Figure S6A). The brain deviations showed positive loadings in the medial occipital, inferior frontal, insula, and dorsal anterior cingulate cortices (dACC), and negative loadings in the dorsal medial and lateral prefrontal and lateral temporal cortices (Figure 3D, Figure S6B). However, no associations between brain deviations and symptoms were observed in the supra-normal subtype.

**Figure 3.**
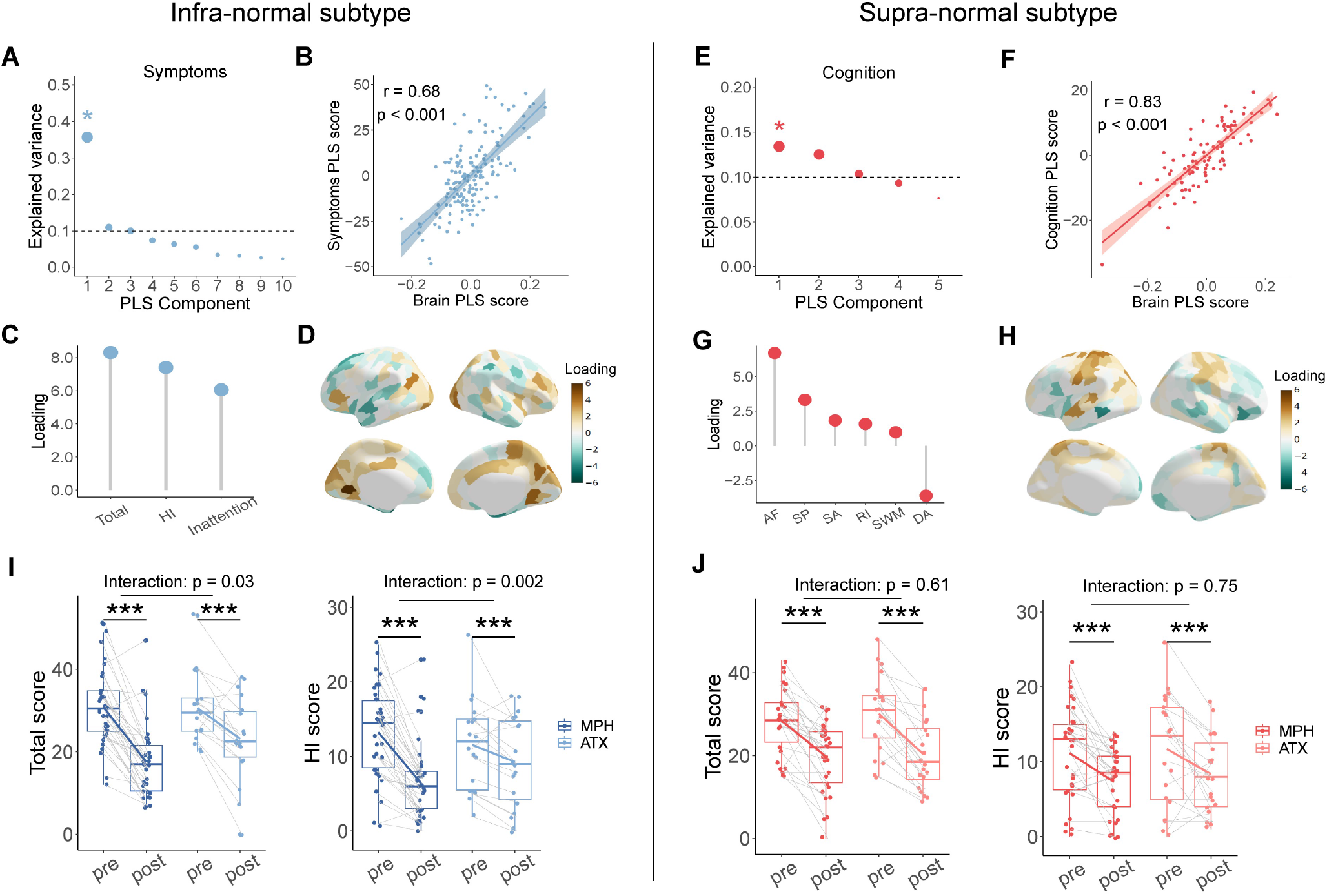
ADHD biotype differences in brain-symptom/cognition relationship and treatment response. **(A-D)** Brain-symptom association in the infra-normal biotype. **(A)** Explained variance for PLS components. The significant PLS component is marked with an asterisk. **(B)** Pearson correlation between brain scores and symptom scores. Shaded areas correspond to 95% CIs. **(C)** Loading values for each symptom variable. **(D)** Loading values for each of the 219 brain regions. **(E-H)** Brain-cognition association in the supra-normal biotype. **(E)** Explained variance for PLS components. The significant PLS component is marked with an asterisk. **(F)** Pearson correlation between brain scores and cognitive scores. Shaded areas correspond to 95% CIs. **(G)** Loading values for each cognitive variable. **(H)** Loading values for each of the 219 brain regions. **(I, J)** Differential treatment response between two ADHD biotypes. **(I)** Significant drug type × treatment interaction in the infra-normal subtype as assessed by total score (left) and HI score (right). **(J)** No drug type × treatment interaction in the supra-normal subtype as assessed by total score (left) and HI score (right). PLS: partial least squares; HI: hyperactivity-impulsivity; AF: attentional flexibility; SP: spatial planning; SA: sustained attention; RI: response inhibition; SWM: spatial working memory; DA: delay aversion; MPH: methylphenidate; ATX: atomoxetine.

In the supra-normal biotype, the first PLS component explained 13.4% of the variance in cognition in ADHD patients (p_permutation_ = 0.006; Figure 3E), and there was a significant correlation between brain scores and cognitive scores (r = 0.83, p_permutation_ < 0.001; Figure 3F). The cognitive contribution loadings showed the highest positive values for attention flexibility, spatial planning, and sustaining attention and negative values for delay aversion (Figure 3G, Figure S6C). Positive loadings of thickness deviation were primarily located in the sensorimotor cortices, while negative loadings were found in the superior frontal, lateral orbitofrontal, and temporal cortices (Figure 3H, Figure S6D). No associations between brain deviations and cognition were observed in the infranormal subtype.

### Differential treatment responses between ADHD subtypes

Pre- and post-treatment symptom scores for each biotype were presented in Table S2. A significant subtype × drug type × treatment interaction was detected for the total symptom score (F = 4.33, p = 0.04; Table S3) and hyperactivity-impulsivity score (F = 3.99, p = 0.048; Table S4) but not for the inattention score (F = 1.93, p = 0.17; Table S5). Post hoc analysis revealed a significant drug type × treatment interaction for the total symptom score (F = 5.25, p = 0.03, Table S6) and hyperactivity-impulsivity score (F = 10.91, p = 0.002, Table S7) in the infranormal subtype (Figure 3I) but not in the supra-normal subtype (Figure 3J). Specifically, the infra-normal subtype responded better to methylphenidate than to atomoxetine as assessed by the total symptom score and hyperactivity-impulsivity score (Figure 3I). For the supra-normal subtype, we observed a main effect of only treatment (F > 24.56, p < 0.001, Figure 3J).

### Subtype-related brain deviations relating to transcriptomic profiles

For the infra-normal subtype, the first PLS component explained 30% of the variance in cortical thickness differences (p_spin_ < 0.001; Figure 4A). There was a positive correlation between the PLS gene score and the spatial pattern of cortical thickness differences (r = 0.55, p_spin_ < 0.001; Figure 4B). The PLS component revealed a transcriptional profile with low gene expression mainly in the dorsal medial and lateral frontal cortices and high expression in the medial occipital cortex, dACC, insula, and lateral temporal cortex (Figure 4C). Furthermore, we performed GO enrichment analysis for the top-ranked PLS+ (1,459 genes) and PLS-genes (1,414 genes), respectively. The PLS+ genes were enriched in “cytosolic ribosomes”, “regulation of cell activation”, “transcription coregulator activity”, and “embryonic morphogenesis” (p_FDR_ < 0.05; Figure 4D, Figure S7A), and the top-ranked PLS-genes were enriched in “axons”, “presynapses”, “postsynapses”, “modulation of chemical synaptic transmission”, and “regulation of transmembrane transport” (p_FDR_ < 0.05; Figure 4E, Figure S7B). We did not find any brain-gene correlations for the supra-normal subtype.

**Figure 4.**
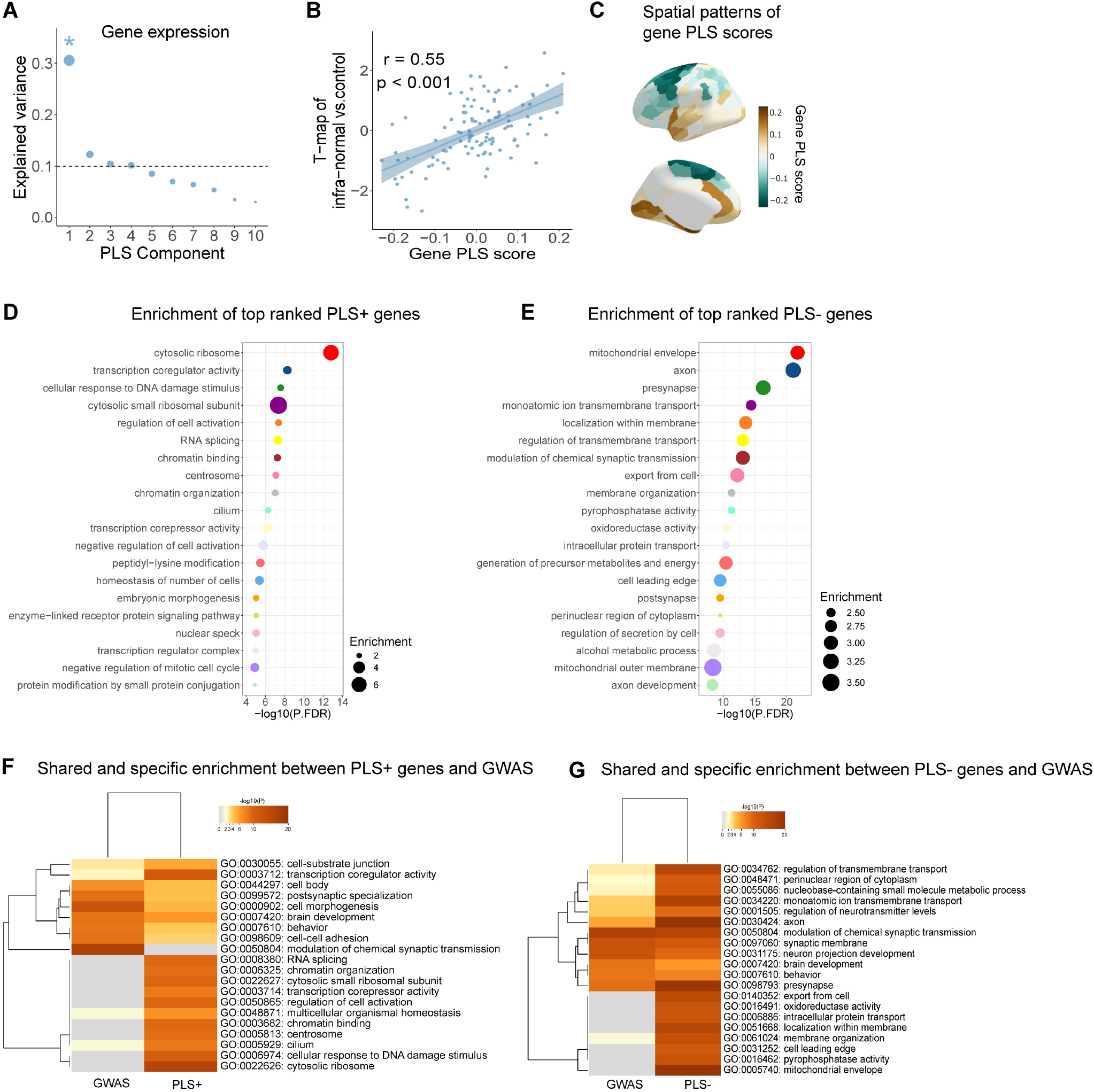
Association between brain alterations of the infra-normal biotype and gene transcriptomic profiles. **(A)** Explained variance for the first 10 components obtained from the PLS analysis. The significant PLS component is marked with an asterisk. **(B)** Pearson correlation between gene scores and *T*-map of the infra-normal biotype. Shaded areas correspond to 95% CIs. **(C)** Spatial patterns of gene PLS scores across 111 brain regions in the left hemisphere. **(D, E)** Functional enrichment for top ranked PLS+ genes **(D)** and PLS-genes **(E)**. Significant GO terms are shown with the node size denoting fold enrichment. **(F, G)** Results of multi-gene-list meta-analysis using PLS genes and polygenic risk for ADHD (F: PLS+ genes, G: PLS-genes). The heatmap shows shared and specific enrichment. Cell color denotes significant p-value and gray color denotes non-significant enrichment.

To further evaluate whether the PLS+ or PLS-genes were enriched in similar terms as the polygenic risk genes for ADHD, a meta-analysis of multiple gene lists was performed. The PLS+ and GWAS genes had common GO terms, including “transcription coregulator activity”, “embryonic morphogenesis”, “regulation of cell activation”, and “transcription regulator complex” (Figure 4F, Figure S8A). The PLS- and GWAS genes shared GO terms, including “axon”, “presynapse”, “modulation of chemical synaptic transmission”, and “regulation of transmembrane transport” (Figure 4G, Figure S8B).

## Discussion

Using normative models of cortical morphology, we identified the neuroanatomical heterogeneity of ADHD and defined two biotypes: an infra-normal subtype with cortical thinning and a supranormal subtype with cortical thickening. The two ADHD biotypes were highly reproducible across independent sites and were characterized by distinct symptomatic, cognitive, and gene expression profiles, as well as treatment responses. These findings advance our understanding of the neurobiological basis underlying the clinical heterogeneity of ADHD and suggest the potential value of data-driven biotyping for personalized treatment.

Previous studies of ADHD have reported small effect sizes for case-control differences in cortical thickness. For instance, a previous ENIGMA study (10) revealed that cortical thinning was restricted to four regions (temporal pole, precentral, fusiform, and parahippocampal gyrus) in children with ADHD, with effect sizes ranging from -0.18 to -0.15. The largest structural MRI study of ADHD based on the Adolescent Brain Cognitive Development (ABCD) data (11) reported no significant case-control differences in cortical thickness. Consistent with the previous study, we found no significant case-control differences in cortical thickness when the data from all children with ADHD were pooled. In contrast to these case-control analyses, we observed individual-level deviations in cortical thickness from the normal references in widespread regions. Moreover, the location of these deviations varied considerably, suggesting remarkable neuroanatomical heterogeneity in ADHD patients. Based on these deviated signatures, we identified two reproducible and clinically meaningful subtypes, highlighting the critical need to discover ADHD biotypes.

The two ADHD biotypes showed opposite deviations in cortical thickness, with negative deviations occurring primarily in the infra-normal biotype and positive deviations occurring primarily in the supra-normal biotype. Several previous studies have reported both cortical thinning in the frontal and lateral occipitotemporal lobes (10) and cortical thickening in the occipital lobe (49, 50), which partly support our findings. Interestingly, the two ADHD biotypes were associated with different symptomatic and cognitive profiles. These finding highlights how different brain structural features shape specific phenotypes. Consistent with the “modernized concept of ADHD” (51), our findings highlight the need to move beyond an understanding of ADHD based solely on “average patient”. It is worth noting that the whole-brain deviation pattern, rather than abnormalities in a single region, was associated with the core symptoms or cognition of each biotype. This finding suggests the potential role of inter-regional structural covariance in the clinical and cognitive phenotypes of ADHD.

One clinical benefit of identifying ADHD biotypes is the advancement of personalized treatment, as medications may have different effects on patients with distinct brain characteristics. This idea is supported by our findings that only patients with the infra-normal subtype had a better treatment response to methylphenidate than to atomoxetine. Several previous studies have suggested that stimulants (i.e., methylphenidate) are more effective than nonstimulants (i.e., atomoxetine) for some children with ADHD (52-54). An intriguing question raised by these results is why this differential effect of methylphenidate and atomoxetine is observed specifically in the infra-normal subtype, which is characterized by overall cortical thinning. One possible explanation lies in the differences in neurobiological mechanisms between the two drugs. Both methylphenidate and atomoxetine increase extracellular synaptic levels of dopamine and norepinephrine in the prefrontal cortex by blocking dopamine and norepinephrine transporters (55). However, methylphenidate also increases catecholamine transmission in the striatum and caudate, whereas the effects of atomoxetine are specific to the prefrontal cortex (56). The reduced cortical thickness observed in the infra-normal subtype suggests disruptions in the top-down regulations between numerous cortices and basal ganglia (57). Consequently, the increase in dopamine and norepinephrine levels in the prefrontal cortex and mesolimbic circuit induced by methylphenidate could lead to superior improvements in the core ADHD symptoms in the infra-normal subtype.

The identified ADHD biotypes revealed specific molecular mechanisms that underlie alterations in brain morphology. We observed associations between brain morphology and gene expression only in the infra-normal subtype, with genes enriched in GO cellular components and biological processes involved in neurodevelopment, including pre- and postsynapses, modulation of chemical synaptic transmission, and axon development. Evidence suggests that disruption of synapses is the most common effect of ADHD-related genetic variants (CDH13, DRD4, and SLC6A3) (58-60). Reduced neuronal and synaptic density may lead to cortical thinning in the infra-normal subtype. The pharmacological effects of stimulants also suggest that neurotransmitter dysregulation contributes to this disorder. Furthermore, observations of abnormalities in cell morphogenesis and axon growth (FOXP2, MEF2C, and SLC6A4) (61) are consistent with the immaturation observed in ADHD. Finally, enrichment analysis of the risk genes reported in ADHD GWASs revealed several shared GO terms, particularly those related to neurodevelopmental processes. These results validate and reinforce the reliability of our findings.

This study has several limitations. First, our ADHD sample was drawn from China. Normative models of cortical morphology were established based on the Chinese TDC population. Given the anatomical differences between Chinese and Caucasian populations (62), future research needs to replicate these findings in samples of diverse geographical origins. Second, we demonstrated subtype-specific medication effects on symptoms. However, how different medications influence brain phenotypes in these two subtypes warrants further investigation. Third, we used the AHBA transcriptomic dataset to establish associations between normally expressed genes and ADHD-related brain phenotypes. The AHBA donors were all adults, whereas our ADHD subjects were children. Future studies should include genes from postmortem brain tissue of individuals with ADHD, allowing for a more direct exploration of the links between MRI-based brain phenotypes and histologically measured dysregulated genes.

## Contributors

XB, QC, LY and YH conceived and designed the study. LY, QC, and YW collaborated in the design, data management, quality control and analysis processes. YZ, ZF, XZ, and KZ collected the data for the PKU6 cohort, LS collected the data for the ADHD200-PKU, and LM, NL, JL, GZ, YD, JW, RC, HZ, WM, YW, JG, ST, SQ, ST, QD, and YH collected the data for the BNU cohort. XB performed the data analysis and produced the figures. MX and ZC checked the accuracy of the data analysis. XB wrote the first draft of the manuscript with critical revision from MX, ZC, and YH. All the authors reviewed the manuscript and agreed with the contents of this study. YH was the project supervisor. XB and YH were responsible for the final decision to submit for publication.

## Data sharing

The ADHD200 dataset is freely available at https://fcon_1000.projects.nitrc.org/indi/adhd200/. The Allen Human Brain Atlas is available at https://human.brain-map.org/static/download. The remaining datasets will be made available from the corresponding author upon reasonable request.

## Acknowledgments

YH was supported by the National Natural Science Foundation of China (grant no. 82021004). QC was supported by the STI 2030—Major Projects (grant no. 2021ZD0200509) and the National Natural Science Foundation of China (grant no. 82371548). LY was supported by the National Natural Science Foundation of China (grant no. 81671358). MX was supported by the Beijing Natural Science Foundation (grant no. JQ23033). ZC was supported by STI 2030-Major Projects (grant no. 2022ZD0211300).

